# Endocannabinoids and dopamine balance basal ganglia output

**DOI:** 10.1101/2020.05.18.099341

**Authors:** Lilach Gorodetski, Yocheved Loewenstern, Anna Faynveitz, Izhar Bar-Gad, Kim T. Blackwell, Alon Korngreen

**Affiliations:** The Leslie and Susan Gonda Interdisciplinary Brain Research Center, Bar-Ilan University, Ramat Gan 52900, Israel; The Mina and Everard Goodman Faculty of Life Sciences, Bar-Ilan University, Ramat Gan 52900, Israel; Department of Bioengineering, George Mason University, Fairfax, Virginia, USA

**Author notes:** Corresponding author: Prof. Alon Korngreen, Bar Ilan University, The Leslie and Susan Gonda Interdisciplinary Brain Research Center, The Mina and Everard Goodman Faculty of Life Sciences, RamatGan 52900, Israel.

**Keywords:** entopeduncular nucleus, endocannabinoids, long-term plasticity, basal ganglia

## Abstract

The entopeduncular nucleus is one of the basal ganglia’s output nuclei, thereby controlling basal ganglia information processing. Entopeduncular nucleus neurons integrate GABAergic inputs from the striatum and the globus pallidus and glutamatergic inputs from the subthalamic nucleus. We show that endocannabinoids and dopamine interact to modulate the long-term plasticity of all the primary afferents to the entopeduncular nucleus. Our results suggest that dopamine-endocannabinoids interplay determines the balance of the direct or indirect dominance of entopeduncular nucleus output. Furthermore, we demonstrate that, despite the lack of axon collaterals, information is transferred between neighboring neurons in the entopeduncular nucleus via endocannabinoids diffusion. These results transform the prevailing view of the entopeduncular nucleus as a feedforward “relay” nucleus to an intricate control unit, which may play a vital role in the process of action selection.

## Introduction

The basal ganglia (BG) are a set of nuclei implicated in addiction (Hiroi et al., 1999), motor control (Yin, 2010), reinforcement learning (Cromwell et al., 2005; Lex and Hauber, 2010; Yin et al., 2005), and many brain disorders such as Parkinson’s disease, Huntington’s disease (Alexander et al., 1990; Feigin et al., 2007; Kloppel et al., 2009; Wichmann and DeLong, 1996) and Tourette syndrome (Leckman et al., 2010). The passage of information through the BG is analogous to a funnel having the striatum (STR) for a mouth, the globus pallidus (GP) as a smaller intermediate structure, and the substantia nigra pars reticulata (SNr) together with the entopeduncular nucleus (EP in rodents, homologues to the GPi in primates) as the nozzle. This funneling is characterized by a reduction in the neuronal population (in the rodent), which goes from millions in the STR to only thousands in the EP, where the direct, indirect, and hyperdirect pathways of the BG converge (Bevan et al., 1997; Bolam and Smith, 1992; Bosch et al., 2012; Nagy et al., 1978; Nambu, 2004; Nambu et al., 2002; Van Der Kooy and Carter, 1981). This structure dictates the basic features of basal ganglia computation and dimensionality reduction (Bar-Gad et al., 2003; Bar-Gad and Bergman, 2001).

Cellular integration of excitatory (Bosch et al., 2012; Nambu, 2004; Nambu et al., 2002) and inhibitory (Smith et al., 1998) inputs from all BG pathways by individual neurons in the EP determines BG output (Bugaysen et al., 2013; Kaneda et al., 2007; Kaneda and Kita, 2005; Kim and Kita, 2013; Kita, 2001; Kita et al., 2004). A vital feature of the EP is the lack of axon collaterals between its neurons, defining it as a pure feedforward nucleus (Parent et al., 2000). In rodents and primates, the direct pathway GABAergic axons from the STR connect mostly to the dendrites of EP neurons distal to the soma, while the synapses of the indirect pathway GP fibers are perisomatic (Bolam and Smith, 1992; Hazrati et al., 1990; Hazrati and Parent, 1992; Smith et al., 1998). In addition to this structural polarization, the EP also displays functional polarity. STR-EP inhibitory synapses display short-term facilitation (Kim and Kita, 2013; Sims et al., 2008), while the GP-EP synapses display short-term depression (Bugaysen et al., 2013; Connelly et al., 2010; Kita, 2001; Sims et al., 2008).

In addition to the interplay between GABAergic inputs from the direct and indirect pathways, the EP receives glutamatergic inputs, which exhibit endocannabinoid receptor (CB1R) dependent LTD (Gorodetski et al., 2018). Interestingly, endocannabinoid (eCB) receptors are essential modulators of both inhibitory and excitatory synaptic transmission (Castillo et al., 2012; Katona and Freund, 2012). This universal function of eCBs led us to hypothesize that endocannabinoids may modulate GABAergic synaptic integration in the EP and that, as in the striatum (Centonze et al., 2007; Freiman et al., 2006; Kreitzer and Malenka, 2005; Maccarrone et al., 2008; Narushima et al., 2007, 2006a, 2006b; Uchigashima et al., 2007; Yin and Lovinger, 2006), dopamine would modulate eCB signaling in the EP. We tested these hypotheses using whole-cell patch-clamp recording in acute brain slices. We show that synaptic integration in the EP depends on eCB release and that dopamine modulates eCB dependent plasticity. Our results suggest that, despite the anatomical lack of axon collaterals, there is a lateral transfer of information within and between neurons in the entopeduncular nucleus. Furthermore, we demonstrate that, despite the lack of axon collaterals, information is transferred between neighboring neurons in the EP via eCB diffusion.

## Materials and Methods

All procedures were approved and supervised by the Institutional Animal Care and Use Committee and followed the National Institutes of Health Guide for the Care and Use of Laboratory Animals and the Bar-Ilan University Guidelines for the Use and Care of Laboratory Animals in Research. This study was approved by the Israel National Committee for Experiments in Laboratory Animals at the Ministry of Health.

### Surgery and stereotaxic viral injections

Five LE-Tg(DRad2-icre)1Ottc rats (RRRC Strain Acquisition Coordinator University of Missouri) (of either sex, 1.5-2 months old) were anesthetized using isoflurane, following by an I.M. injection of ketamine HCI (100 mg/kg) and xzylazine HCl (10 mg/kg). The rat’s head was fixed in a stereotaxic frame, and the AAV5-EF1α-DIO-ChR2(H134R)-YFP virus (1µl; University of North Carolina Gene Therapy Center) was injected bilaterally into the GP (AP, −0.95 mm; ML, ± 3 mm; DV, 5.75 mm) (Paxinos and Watson, 2007). The virus was injected using a syringe pump (World Precision Instruments) at a rate of 0.1 µl/min and left in place for 10 minutes after injection to allow viral particle diffusion before the needle removal. Whole-cell experiments were carried out three weeks after viral injection.

### In vitro slice preparation

Brain slices were obtained from Wistar (P15-P22 of either sex) rats as previously described (Lavian and Korngreen, 2016). Rats were anesthetized by Isoflurane and killed by rapid decapitation. The brain was quickly removed and placed in ice-cold artificial cerebrospinal fluid (ACSF) containing (in mM): 125 NaCl, 2.5 KCl, 15 NaHCO_3_, 1.25 Na_2_HPO_4_, 2 CaCl_2_, 1 MgCl_2_, 25 glucose and 0.5 Na-ascorbate (pH 7.4 with 95% O_2_/5%CO_2_). In all experiments, the ACSF contained APV (50 μM) and CNQX (15 μM) to block NMDA, AMPA receptors, or GABAzine (20 μM) to block GABA receptors. In some experiments, we also added AM251 (3 μm) to block CB1 receptors, Sulpiride (3 μM) to block D2 receptors (D2R), R-SCH23390 (10 μM) to block D1 receptors (D1R), or Quinpirole (5 μM) the D2R agonist. AM251 was dissolved in Dimethyl sulfoxide (DMSO). The final concentration of DMSO was 0.15%. Thick sagittal slices (320–350 μm) were cut at an angle of 17° to preserve functional connectivity between the STN and EP to the midline on an HM 650 V Slicer (Microm International, Walldorf, Germany) and transferred to a submersion-type chamber where they were maintained for the remainder of the day in ACSF at room temperature. Experiments were carried out at 37°C, and the recording chamber was constantly perfused with oxygenated ACSF.

Optogenetic experiments in brain slices were performed on 60-90 days old Long-Evans LE-Tg (Drad2-icre)1Ottc rats 3 weeks following viral injection. To prepare brain slices, rats were deeply anesthetized using ketamine (100mg/kg) and xylazine (10mg/kg) and perfused transcardially with Ice-cold N-Methyl-D-glucamin (NMDG)–ACSF containing (in mM): 92 NMDG, 2.5 KCl, 30 NaHCO3, 1.25 Na2HPO4, 0.5 CaCl2, 10 MgSO4*7H2O, 20 HEPES, 25 glucose, 2 thiourea and 5 Na-ascorbate (pH 7.4). The brain was quickly removed and place in the Ice-cold NMDG-ACSF. Sagittal slices were cut as described above and transferred to chamber contain HEPES holding ACSF containing (in mM): 92 NaCl, 2.5 KCl, 1.25 NaH_2_PO_4_, 30 NaHCO_3_, 20 HEPES, 25 glucose, 2 thiourea, 5 Na-ascorbate, 3 Na-pyruvate, 2 CaCl_2_ and 2 MgSO_4_ (pH 7.4 with 95% O_2_/5%CO_2_).

### In vitro electrophysiology

Individual EP neurons were visualized using infrared differential interference contrast microscopy using an Olympus BX51WI microscope with a 60x water immersion objective (Gorodetski et al., 2018; Lavian et al., 2017; Lavian and Korngreen, 2016). Electrophysiological recordings were performed in the whole-cell configuration of the patch-clamp technique under visual control using a CCD camera (Retiga-Electro, QImaging). Recordings were obtained from the soma of EP neurons using patch pipettes (4–8MΩ) pulled from thick-walled borosilicate glass capillaries (2.0 mm outer diameter, 0.5 mm wall thickness, Hilgenberg, Malsfeld, Germany). The standard pipette solution contained (in mM): 140 K-gluconate, 10 NaCl, 10 HEPES, 4 MgATP, 0.05 Spermin, 5 L-glutathione, 0.5 EGTA and 0.4 GTP (pH 7.2 with KOH; Sigma, St Louis, MO, USA). Under these conditions, the Nernst equilibrium potential for chloride was calculated to be –69 mV. The reference electrode was an Ag–AgCl pellet placed in the bath. Voltage signals were amplified by an Axopatch-200B amplifier or Axopatch-700B (Axon Instruments, Union City, CA, USA), filtered at 5 kHz and sampled at 20 kHz. The 10-mV liquid junction potential measured under these ionic conditions was not corrected.

Excitatory and inhibitory synaptic potentials were evoked via a monopolar 2-3 KΩ Narylene-coated stainless-steel stimulating microelectrode positioned in the STN, GP, or Striatum. The stimulation pulse consisted of 100-800 μA biphasic currents (200 μs cathodal followed by 200 μs anodal phase). For the optogenetic experiments, inhibitory synaptic potentials were evoked via ChR2 activation of GP neurons by optical blue led light (473 nm) stimulation consisted of 1-5 ms light pulses (Prizmatix). The input resistance was monitored throughout the experiment. Data were excluded when the input resistance was not stable for the entire experiment.

### Computational modeling

We created a multi-compartment, multi-ion channel model of EP neurons to investigate synaptic integration in response to in vivo like inputs. The model was implemented in the simulator MOOSE, using the moose_nerp python package, which allows declarative model specification (https://github.com/neurord/moose_nerp/tree/master/moose_nerp/ep) and simulated with a timestep of 0.001 ms.

The model includes fast and slow sodium currents, a fast and slow transient potassium current, Kv2 (non-inactivating), and Kv3 (inactivating) potassium channels, small conductance and big conductance calcium-activated potassium channels, two hyperpolarization-activated cyclic-nucleotide gated (HCN) channels, and one high voltage-activated calcium channel. NMDA and AMPA synaptic channels and GABA synaptic channels were distributed along the dendrites. Intracellular calcium concentration was increased by influx through NMDA and calcium channels and decayed with a single time constant. Channel conductances, time constants and voltage dependence of gates, as well as membrane resistivity, axial resistivity, and capacitivity, were adjusted using the automatic parameter optimization algorithm, ajustador (available from https://github.com/neurord/ajustador), to match in vitro EP neuron response to hyperpolarizing and depolarizing current injection (<u>ep032117_2_Waves</u> available from https://github.com/neurord/waves/tree/master/EPmeasurements).

Synaptic inputs were created with a mean ISI and coefficient of variation of ISI similar to that measured in vivo (GPe: 29 Hz, (Kita and Kita, 2011), STR: 4 Hz (for computational efficiency, each input train represents four trains firing at 1 Hz) (Kita and Kita, 2011), STN: 18 Hz, (Wilson and Bevan, 2011)). To measure information processing, these mean firing rates were modulated with a different frequency for each type of input (inhomogeneous Poisson process). The oscillation frequencies differed by a factor of ∼3 to prevent harmonics overlapping the main frequencies. The code for spike trains is available from https://github.com/neurord/synth_trains/. STR synaptic inputs were distributed along the dendrites, GPe inputs distributed within 60 µm of the soma, and STN inputs everywhere. Independent trials were created by selecting a different subset of the set of spike trains and randomly selecting the location of the target synapse. Model output analysis was conducted using python 3.6; power spectra of the model output were calculated using the FFT function in NumPy, and then averaged across the set of trials.

### Analysis and statistics

All off-line analyses of experimental data were carried out using IgorPro 7.0 (WaveMetrics; RRID:SCR_000325) and Matlab R2013a (MathWorks; RRID:SCR_001622). The results for each experiment were obtained from at least three rats. The results were pooled and displayed as means±SEM. The steady-state level of LTD and LTP was calculated as the average EPSP or IPSP amplitude 30 min after the depolarization protocols and was presented as the percentage of the average of the baseline (the first 5 min of baseline) EPSP or IPSP amplitude. A Mann-Whitney U test was used to test for significance in all the experiments (in figures p values are denoted as * p<0.05, ** p<0.01 and *** p<0.001).

## Results

### eCBs mediate long term changes in the basal ganglia pathway

We recently showed that post-synaptic depolarization of neurons in the EP induces the release of endocannabinoids, which leads to long-term depression of glutamatergic input to these neurons (Gorodetski et al., 2018). Given the high density of CB1 receptors in the EP (Herkenham et al., 1990), we hypothesized that endocannabinoids might participate in long-term changes to the plasticity of other synaptic inputs to the EP. To investigate this hypothesis, we tested the effect of eCB release on GABAergic inputs to the EP from the direct and indirect pathways of the BG. We carried out whole-cell recordings of the membrane potential from neurons in the EP, while extracellularly stimulating in the GP using a tungsten microelectrode. In the presence of CNQX and APV, brief electrical stimulation to the GP generated inhibitory synaptic responses in the EP (Fig. 1A insert). We only studied synapses displaying clear short-term depression, identifying them as GP-EP synapses (Lavian and Korngreen, 2016). After 5 minutes of stable recording, a 10-second train at 100 Hz of current pulses was injected via the whole-cell electrode to the soma of the EP neuron. This high-frequency post-synaptic stimulation protocol (post-HFS) induced robust LTD (54%±1%, N=10, Figure 1A, p=10^−5^) that was blocked by AM-251 (103%±4%, N=12, Figure 1A, p=0.001). Repeating the experiment with post-HFS induction of 50 Hz resulted in a smaller LTD (74%±2%, N=9, Figure 1B, p=0.02), whereas post-HFS using an induction frequency of 10 Hz did not affect synaptic strength (99%±3%, N=4, Figure 1C, p=0.45). The steady-state level of LTD displayed a monotonic dependence on the frequency of the induction protocol (Fig. 1D) that was similar to the monotonic dependence of the intracellular calcium concentration on firing frequency in the EP (Gorodetski et al., 2018). We did not observe a change in the paired-pulse ratio (PPR) of evoked IPSCs between before and after the post-HFS protocol (N=9, p=0.33).

**Figure 1:**
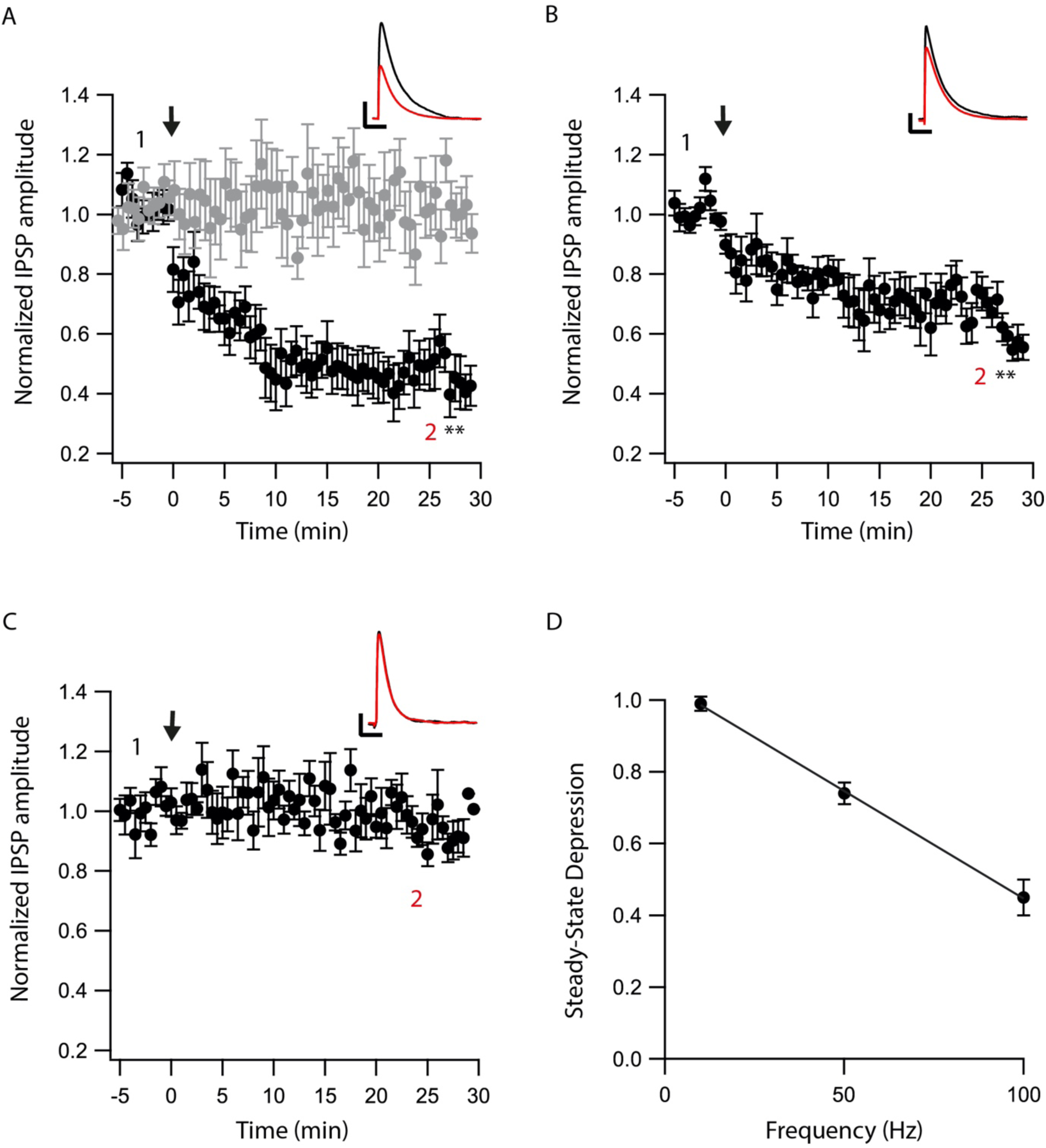
eCBs induce LTD at GP-EP synapses. A. Post-HFS stimulation of 100 Hz for 10 s induced LTD at GP-EP synapses (n=10). Bath application of AM251 blocked the depression at GP-EP synapses (n=13) (grey). Normalized IPSP amplitudes (normalization to the mean amplitude of EPSP recorded during the baseline recordings) are plotted against time. Arrow indicates the time of induction of the post-HFS protocol, and error bars represent the SEM. Inset, traces from a representative experiment illustrating the average IPSP from the first 5 min of the baseline (1) and the last 5 min after stimulation (2). Vertical scale bar is 1 mV and the horizontal scale bar is 50 ms. B. Similar to A only with a 50 Hz post-HFS stimulation (n=9). C. Similar to A only with a 10 Hz post-HFS stimulation (n=4). D. Changes in the steady-state levels of IPSP amplitudes vs. the frequency of the post-HFS protocol presented as the mean±SEM. (** p<0.01)

Next, we tested whether eCBs modulated direct pathway input to the EP. We carried out whole-cell recordings of the membrane potential from EP neurons while applying electrical stimulation to the striatum. In the presence of CNQX and APV, brief electrical stimulation to the striatum generated inhibitory synaptic responses in EP neurons (Fig. 2A insert). Only synapses displaying short-term facilitation, identifying them as STR-EP synapses (Lavian and Korngreen, 2016), were investigated further. After 5 minutes of stable recording, we stimulated the neuron with a 100 Hz post-HFS for 10 s. Contrary to the GP-EP synapse, this protocol induced LTP of the direct pathway input to the EP (144%±11%, N=7, Figure 2A, p=0.0044) that was blocked by AM251 (89%±9%, N=12, Figure 2A, p=0.00001). Contrary to the GP-EP synapse effect, a 50 Hz post-HFS protocol generated almost no change in synaptic plasticity (107%±6%, N=8, p=0.55, Figure 2B). Finally, a 10 Hz post-HFS protocol had no long-term effect on synaptic plasticity in STR-EP synapses (99%±4%, N=6, Figure 2C, p=0.58), similar to the GP-EP synapses. Again, we did not observe a change in the paired-pulse ratio (PPR) of evoked IPSCs between before and after the post-HFS protocol (N=4, p=0.66). This non-linear dependence of LTP on post-synaptic firing agreed with the partial invasion of back-propagating APs into the dendritic tree of neurons in the EP leading to dendritic calcium transients only at high firing rates (Gorodetski et al., 2018).

**Figure 2:**
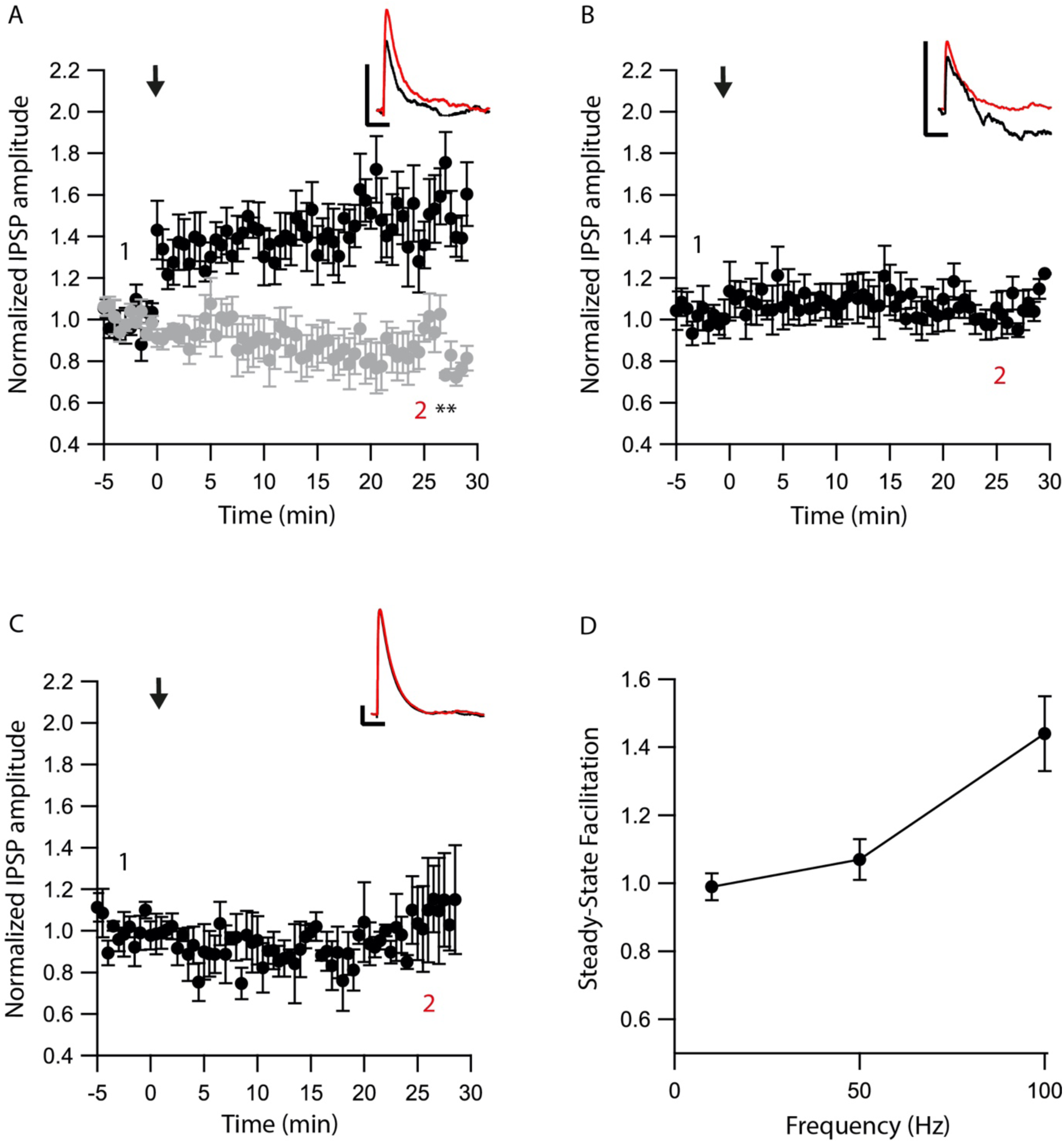
eCBs induce LTP at STR-EP synapses. A. Post-HFS stimulation of 100 Hz for 10 s induced LTP at Str-EP synapses (n=7) compared to the baseline response. Bath application of AM251 blocked the depression at GP-EP synapses (n=12) (grey). Normalized IPSP amplitudes (normalization to the mean amplitude of EPSP recorded during the baseline recordings) are plotted against time. Arrow indicates the time of induction of the post-HFS protocol, and error bars represent the SEM. Inset, traces from a representative experiment illustrating the average IPSP from the first 5 min of the baseline (1) and the last 5 min after stimulation (2). Vertical scale bar is 1 mV and the horizontal scale bar is 50 ms. B. Similar to A only with a 50 Hz post-HFS stimulation (n=4). C. Similar to A only with a 10 Hz post-HFS stimulation (n=6). D. Changes in the steady-state levels of IPSP amplitudes vs. the frequency of the post-HFS protocol presented as the mean±SEM. (** p<0.001)

While we previously showed that post-synaptic depolarization induces eCB dependent synaptic plasticity of glutamatergic input to the EP (Gorodetski et al., 2018), the frequency dependence of this plasticity is unknown. Thus, we applied similar post-HFS protocols as described above while stimulating extracellularly in the STN in the presence of gabazine. Similarly to the results obtained in the GP-EP synapse (Fig. 1), a post-HFS protocol at 100Hz generated robust LTD (60%±10%, N=8, Figure 3A, p=0.04) that was blocked by AM251 (98%±3%, Fig. 3A, p=0.0002), post-HFS at 50 Hz generated a smaller LTD (83%±2%, N=13, Figure 3B, p=0.002) and post-HFS at 10 Hz APs did not generate significant changes in the synaptic plasticity (101%±7%, N=9, Figure 3C, p=0.9145). The changes to the glutamatergic synaptic plasticity were linearly dependent on the frequency of the post-synaptic stimulation (Fig. 4D). Furthermore, we did not observe a change in the paired-pulse ratio (PPR) of evoked EPSCs between before and after the post-HFS protocol (N=6, p=0.08).

**Figure 3:**
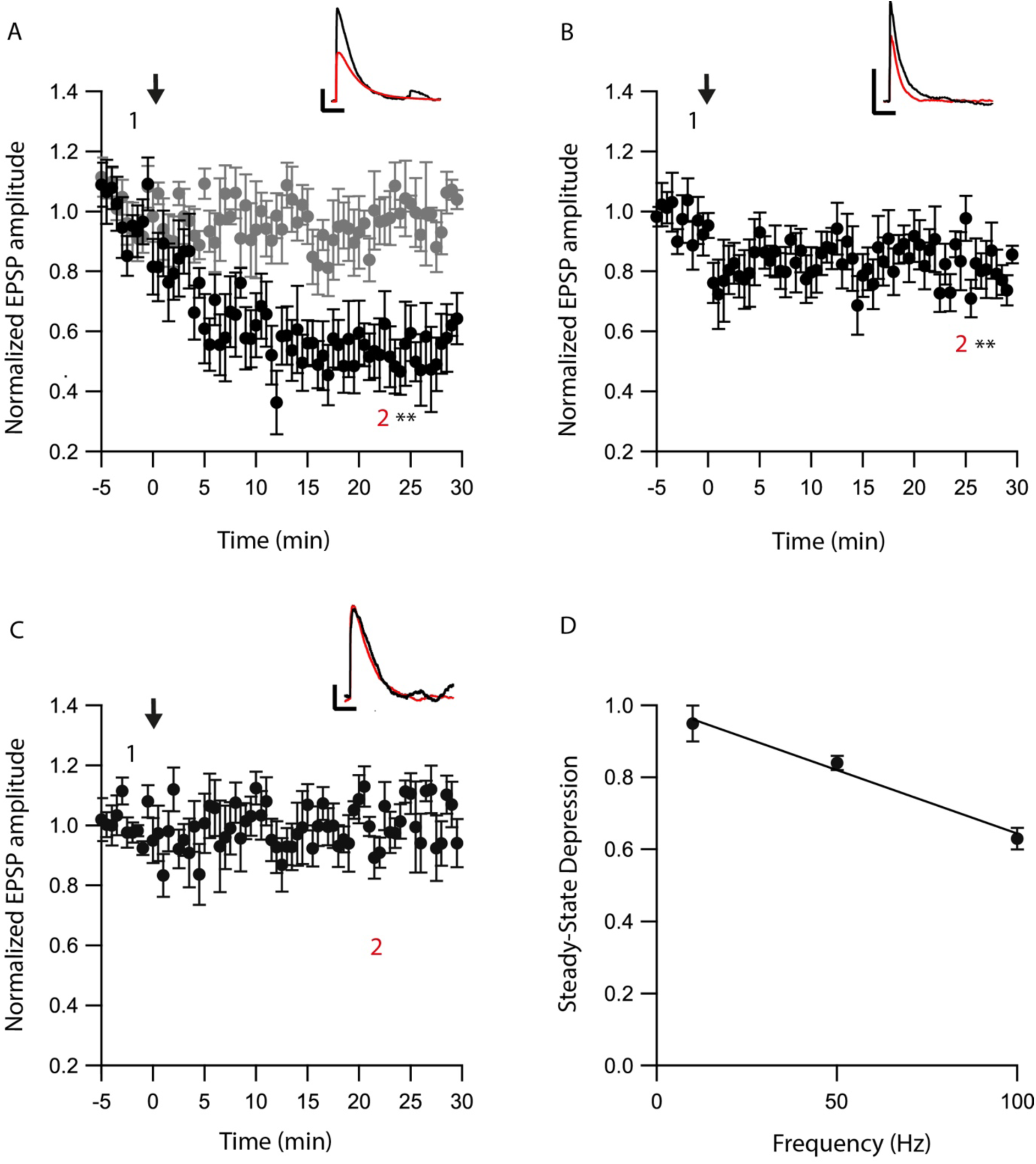
eCB induce LTD at STN-EP synapses. A. Post-HFS stimulation of 100 Hz for 10 s induced LTD at STN-EP synapses (n=8) compared to the baseline response. Bath application of AM251 blocked the depression at STN-EP synapses (n=10) (grey). Normalized EPSP amplitudes (normalization to the mean amplitude of EPSP recorded during the baseline recordings) are plotted against time. Arrow indicates the time of the post-HFS protocol, and error bars represent the SEM. Inset, traces from a representative experiment illustrating the average EPSP from the first 5 min of the baseline (1) and the last 5 min after stimulation (2). Vertical scale bar is 1 mV and the horizontal scale bar is 50 ms. B. Same as in A only with a 50 Hz post-HFS stimulation (n=13). C. Same as in A only with a 10 Hz post-HFS stimulation (n=9). D. Changes in the EPSPs amplitude vs. the frequency of the depolarization of the EP neurons presented as the mean±SEM. (** p<0.01).

**Figure 4:**
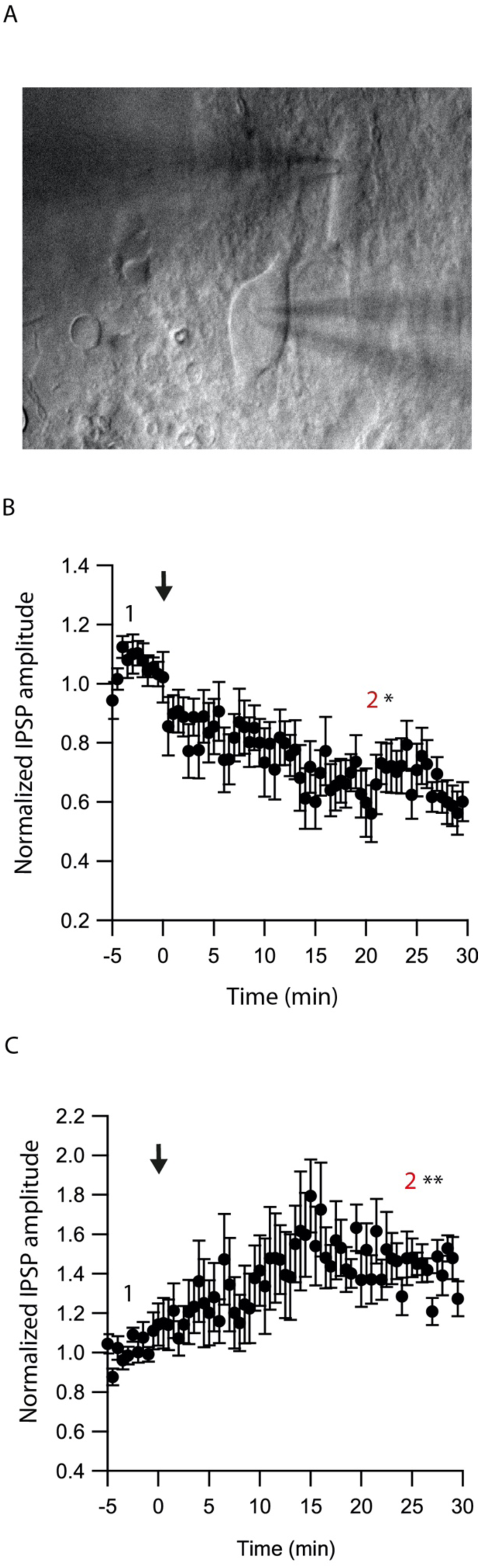
eCBs induce synaptic plasticity in neighboring neurons. A. an image of a paired recording from two neurons in the EP. B. Post-HFS at 100 Hz for 10 s to one neuron induced LTD at GP-EP synapses of the non-stimulated neuron (n=13). Normalized IPSP amplitudes (normalization to the mean amplitude of IPSP recorded during the baseline recordings) are plotted against time. Arrow indicates the time of induction of the depolarization protocol, and error bars represent the SEM. C. Post-HFS at 100 Hz for 10 s to one neuron induced LTP at STR-EP synapses of the non-stimulated neuron (n=5).

### eCBs modulate plasticity in neighboring neurons

As retrograde messengers, eCBs diffuse and affect synapses in surrounding neurons (Kreitzer and Regehr, 2001; Maejima et al., 2001; Ohno-Shosaku et al., 2001; Yanovsky et al., 2003; Zhu, 2005). To test whether this lateral transfer of information occurs in the EP, we performed paired recordings from neighboring neurons (inter-somatic distance < 40 µm) in the EP (Fig. 4A) and measured eCB induced plasticity (Fig. 4). We placed an extracellular stimulating electrode in the GP and applied a single stimulus to test for a synaptic connection to at least one of the neurons. We stimulated the soma of one of the neurons with a 10 s post-HFS at 100 Hz and measured changes in synaptic strength of inputs to the unstimulated neuron (Fig. 4B).

Post-HFS of one neuron generated LTD in the GP-EP GABAergic synapse on the unstimulated neuron (73%±8%, N=13, Figure 4B, p=0.0012). Interestingly, LTD kinetics were slower in neighboring neurons (Fig. 4B) than in the stimulated neurons (Fig. 1A), which may be due to eCB diffusion between neurons. Because concentration decreases with distance for diffusing molecules, such a mechanism suggests that the LTD would be lower in more distant neurons; however, we did not detect a correlation between the magnitude of LTD and the distance between the two somata. Placing the stimulating electrode in the STR produced a similar trend: eCB release by post-HFS in one neuron generated LTP of the STR-EP GABAergic synapse on a neighboring neuron (138%±15%, N=8, Figure 4C, p=0.004). As in the case of the GP-EP synapse, LTP kinetics was slower than that observed in single neurons (Fig. 2A).

### Dopamine modulates eCB induced plasticity

We have shown that dopamine modulates GABAergic inputs to the EP (Lavian et al., 2017). D1Rs modulate GABAergic input from the STR to the EP (Lavian et al., 2017), whereas D2Rs modulate GABAergic input from the GP to the EP. As dopamine interacts with eCBs in other regions of the basal ganglia, we tested the hypothesis that dopamine modulates eCB induced plasticity in the EP. We repeated the experiments described in figures 1-3 in the presence of dopamine antagonists. We measured the change in STR-EP synapses caused by post-HFS (repeating the experiments shown in figure 2A) in the presence of R-SCH23390 (10 μM), the D1R antagonist. Under these conditions, post-HFS for 10 s at 100 Hz did not generate LTP (96%±6%, N=5, Figure 5A, p=0.008) compared to the response of STR-EP synapses (Fig. 2A). Then, we measured the change in GP-EP synapses caused by post-HFS (repeating the experiment shown in Fig. 1A) in the presence of the D2R antagonist, Sulpiride (3 μM). These conditions resulted in LTP of the GP-EP synapse (137%±5%, N=7, Figure 5B, p= 0.002) instead of LTD (Fig. 1A). It has been shown that D2Rs modulate glutamatergic input to the EP. Indeed, 3 µM of Sulpiride blocked the LTD (103%±12%, N=10, figure 5C, p=0.002) that we observed in the experiment without the blocker (Fig. 3A).

**Figure 5:**
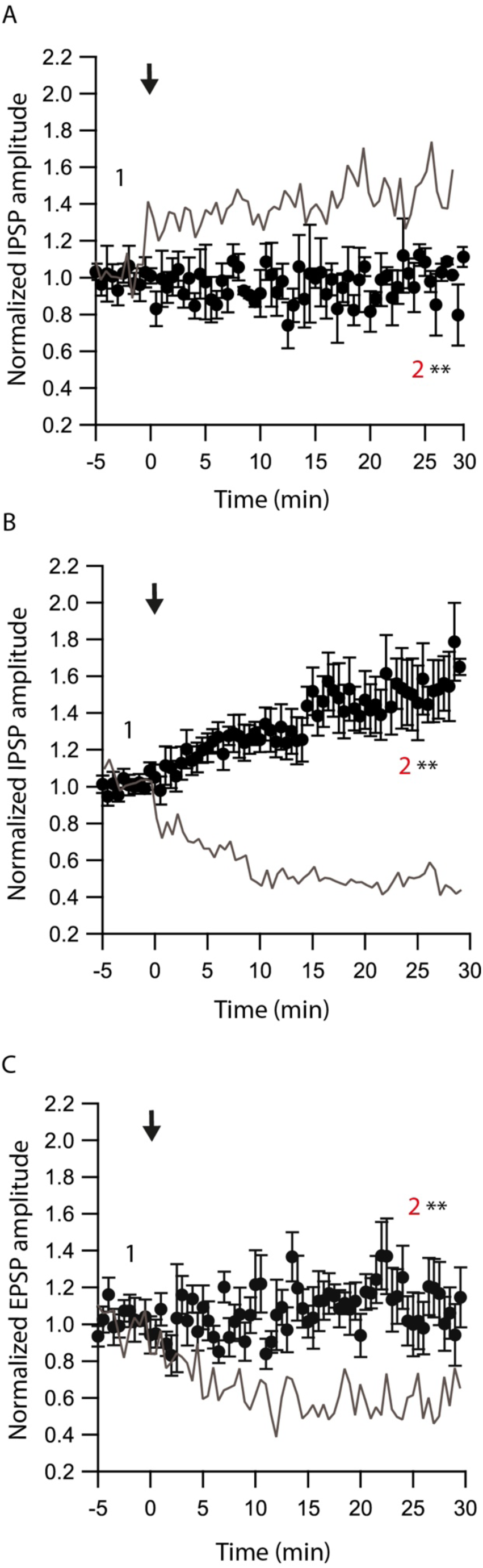
Dopamine modulates eCB induced plasticity. A. Post-HFS stimulation of 100 Hz for 10 s in the presence of D1 antagonist at STR-EP synapse (n=4) compared to control condition from figure 2 in grey. Normalized IPSP amplitudes (normalization to the mean amplitude of IPSP recorded during the baseline recordings) are plotted against time. Arrow indicates the time of induction of the depolarization protocol, and error bars represent the SEM B. Post-HFS stimulation of 100 Hz for 10 s generated LTP in the presence of a D2 antagonist at the GP-EP synapses (n=7) compared to control condition from figure 1 in grey. C. Post-HFS stimulation of 100 Hz for 10 s in the presence of a D2 antagonist at STN-EP synapses (n=10) compared to the control condition from figure 3 in grey (** p<0.001).

The experiments presented in figure 5 demonstrate that dopamine modulates eCB induced plasticity in the EP. However, the source of dopamine is unclear. In slice experiments, dopamine release in the EP can result from spontaneous firing of dopaminergic neurons, or nonspecific electrical stimulation generated action potentials in dopaminergic axons in upstream BG regions. To differentiate between these possibilities, we performed additional experiments using optogenetics. We injected adeno-associated virus (AVV) encoding a fusion of channelrhodopsin-2 and enhanced yellow fluorescent protein (ChR2-YFP) into the GP of LE-Tg (DRad2-icre) rats (Fig. 6Ai). We observed ChR2-YFP expressing somata in the GP (figure 6Aii) but only axonal projections in the EP (Fig. 6Aiii) confirming the localization and specificity of the AAV infection. The GP neuron’s firing was locked to individual light pulses within a stimulation train, confirming ChR2-YFP expression in the cell body (Fig. 6B). Next, we optogenetically stimulated neurons in the GP while performing whole-cell recordings in the EP. In the presence of CNQX and APV, brief optical stimulation to the GP generated inhibitory synaptic responses in EP neurons. Following a 5 minute control period, a 10 s post-HFS at 100 Hz was applied to the soma of the neuron in the EP generating LTP (148%±23%, N=6, figure 6C, p=0.0770) – similar to the LTP produced with electrical stimulation and D2R blocked, thus mirroring the effect observed using electrical stimulation. Bath application of the D2R agonist, Quinpirole (5 μM), produced a small LTD (83%±12%, N=4, figure 6D, p=0.02). These experiments suggest that in slice experiments, in addition to the release of GABA, electrical stimulation of the GP produced dopamine release from dopaminergic axons.

**Figure 6:**
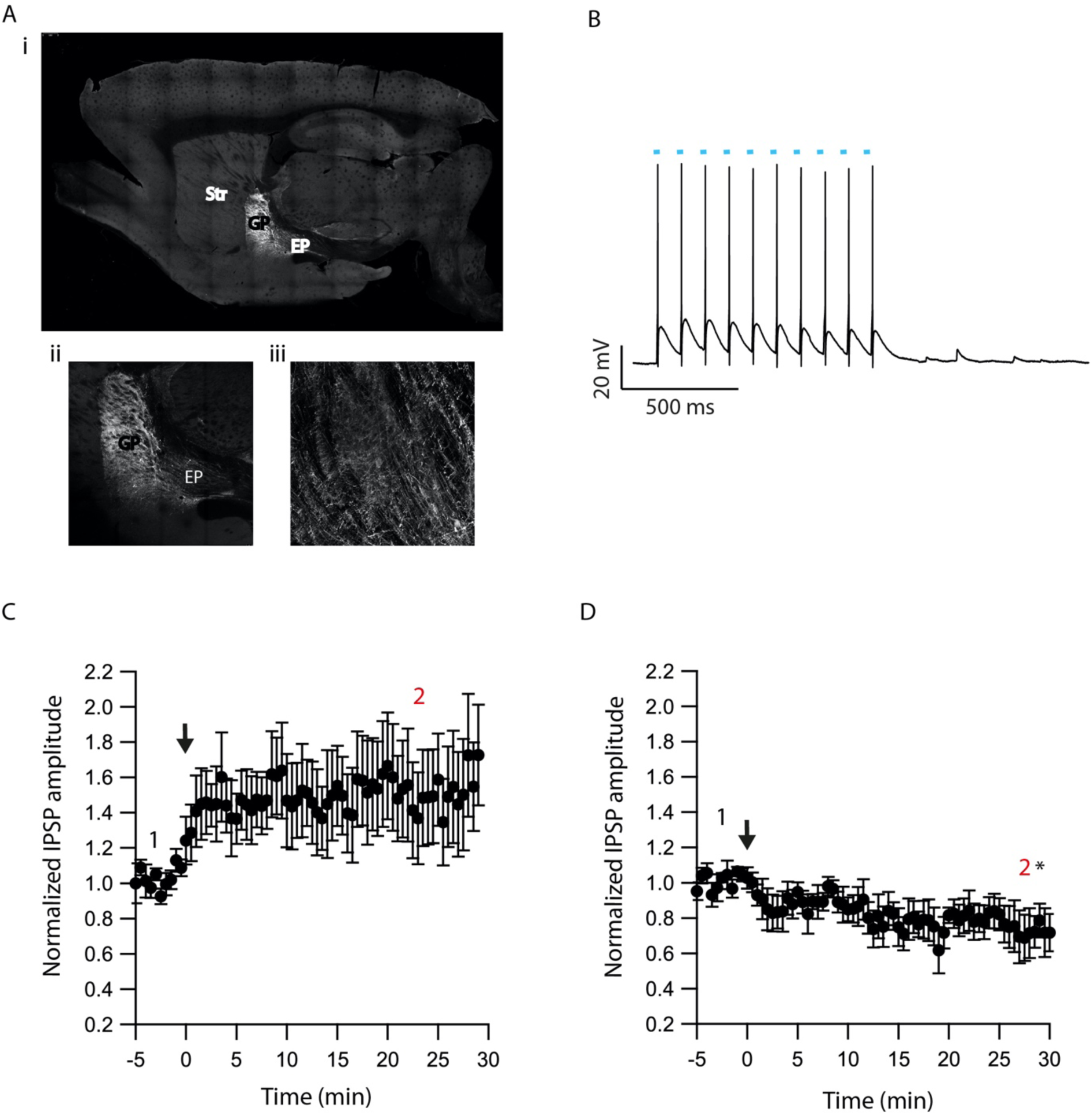
Optogenetic verification of eCB induced plasticity. A. Sagittal brain slice image containing the GP (the injected zone), the EP (the recording zone) and adjacent structures. ChR2-YFP expression was observed in infected cells in the GP and in axons terminals in the EP. The Image was magnified x10 (i), x20 (ii,iii) B. Whole-cell recording from a ChR2 expressing GP neuron illuminated with a 10 Hz light pulse train. The blue dots indicate illumination periods. C. Post-HFS stimulation of 100 Hz for 10 s induced LTP at GP-EP synapses (n=6). Normalized IPSP amplitudes (normalization to the mean amplitude of IPSP recorded during the beginning of baseline recordings) are plotted against time. The arrow indicates the time of induction depolarization protocol, and error bars represent the SEM. D. Post-HFS stimulation of 100 Hz for 10 s generated mild LTD in the presence of a D2 agonist at GP-EP synapses (n=4).

### Dopamine and eCBs modulate EP firing

To demonstrate the role of eCB dependent synaptic plasticity in information processing, we performed simulations of EP neuron activity in response to simultaneous STN, GPe, and STR synaptic inputs. We created a multi-compartmental EP neuron model (Fig 7) by using automatic parameter optimization to adjust the conductance of fast and slow sodium currents, a fast and slow transient potassium current, Kv2 (non-inactivating), and Kv3 (inactivating) potassium channels, small conductance, and big conductance calcium-activated potassium channels, two hyperpolarization-activated cyclic-nucleotide gated (HCN) channels, and one high voltage-activated calcium channel. Fig 7A shows the spontaneous firing of the EP neuron and the typical sag in response to hyperpolarizing current injection. The frequency-current injection curves (Fig 7B) and calcium concentration (Fig 7C) matched experimental data. Using this data-driven model, we assessed the effect of short-term plasticity of STR or GPe inputs to the EP. We performed simulations using STP equations derived from fitting experimental data (Lavian and Korngreen, 2019, 2016). Then we simulated the response to 20 Hz stimulation in the presence of log-normally distributed GPe (29 Hz), SPN (4 Hz – to efficiently model the massive convergence from STR to EP, each input train represents four trains firing at 1 Hz) and STN input (18 Hz), both with and without STP of the 20 Hz inputs (STP was always present for the log-normal inputs to maintain ∼20 Hz EP neuron firing frequency). First, we measured the effect of STP on EP firing frequency. Fig 8A1 shows that 20 Hz GPe inputs produce a 25% reduction in EP firing (No STP) but that this reduction is weaker and more transient with STP. In contrast, Fig 8A2 shows that without STP, STR inputs have a non-significant effect on EP neuron firing, but that with STP the effect of STR inhibition increases, with 50% decrease in EP firing after 300 ms. We further evaluated phase locking of the EP neurons to the 20 Hz input by plotting the power spectral density (PSD). Fig 8B shows a peak in the PSD at 20 Hz, representing regular 20 Hz firing in the presence of the 20 Hz input (compared to basal – the absence of 20 Hz input). The peak at 20 Hz is greater in the absence of STP for GPe inputs, and in the presence of STP for STR inputs.

**Figure 7:**
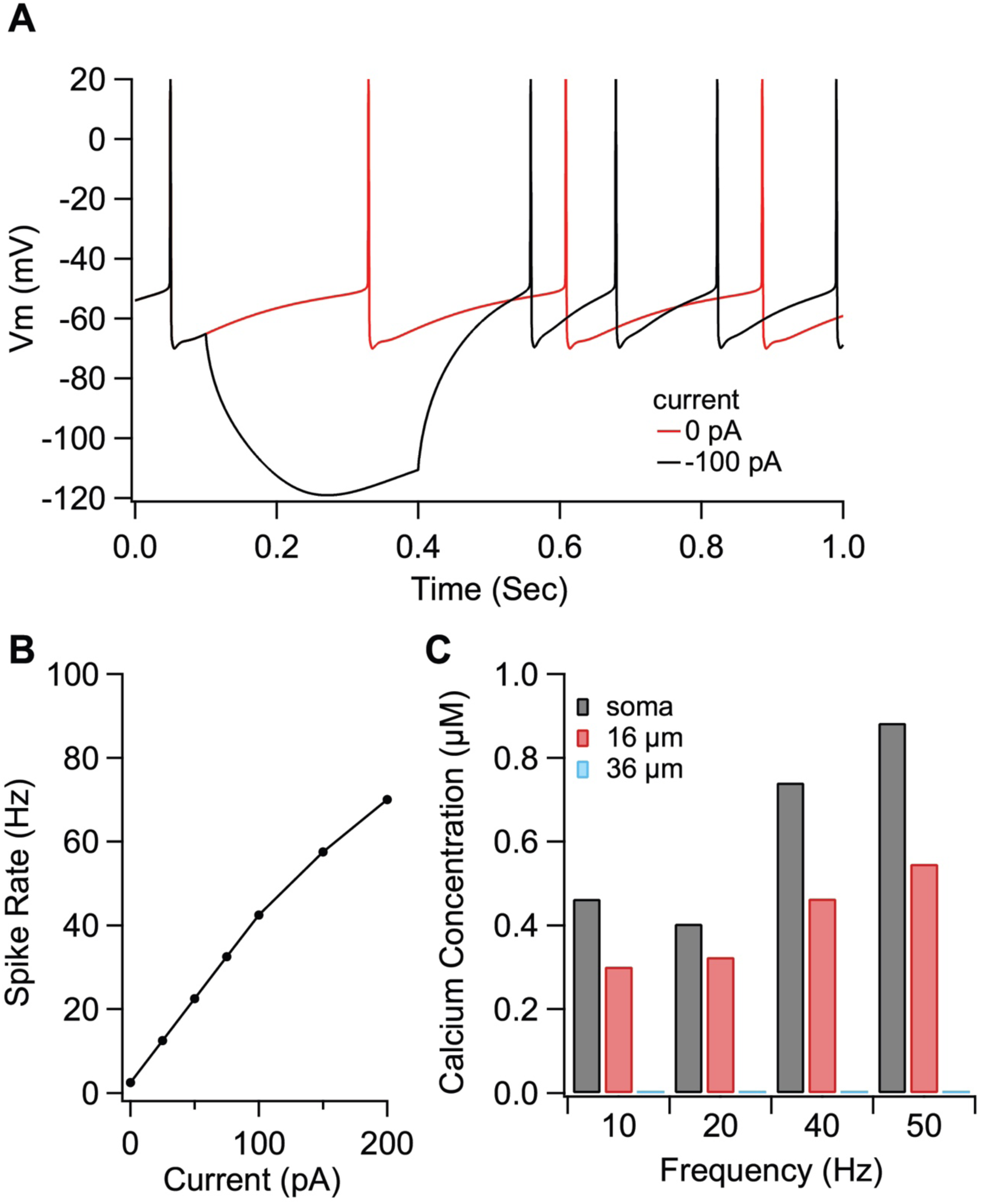
Optimized compartmental model for a neuron in the EP. (A) Parameter optimization to EP neuron data produces multi-compartmental EP neuron model with realistic spiking and rectification characteristics. (A) sample traces of EP neuron model showing spontaneous activity and response to hyerpolarizing current injection. (B) firing frequency versus current injection is similar to that measured experimentally. (C) Calcium concentration versus distance from soma and firing frequency is similar to that measured experimentally.

**Figure 8:**
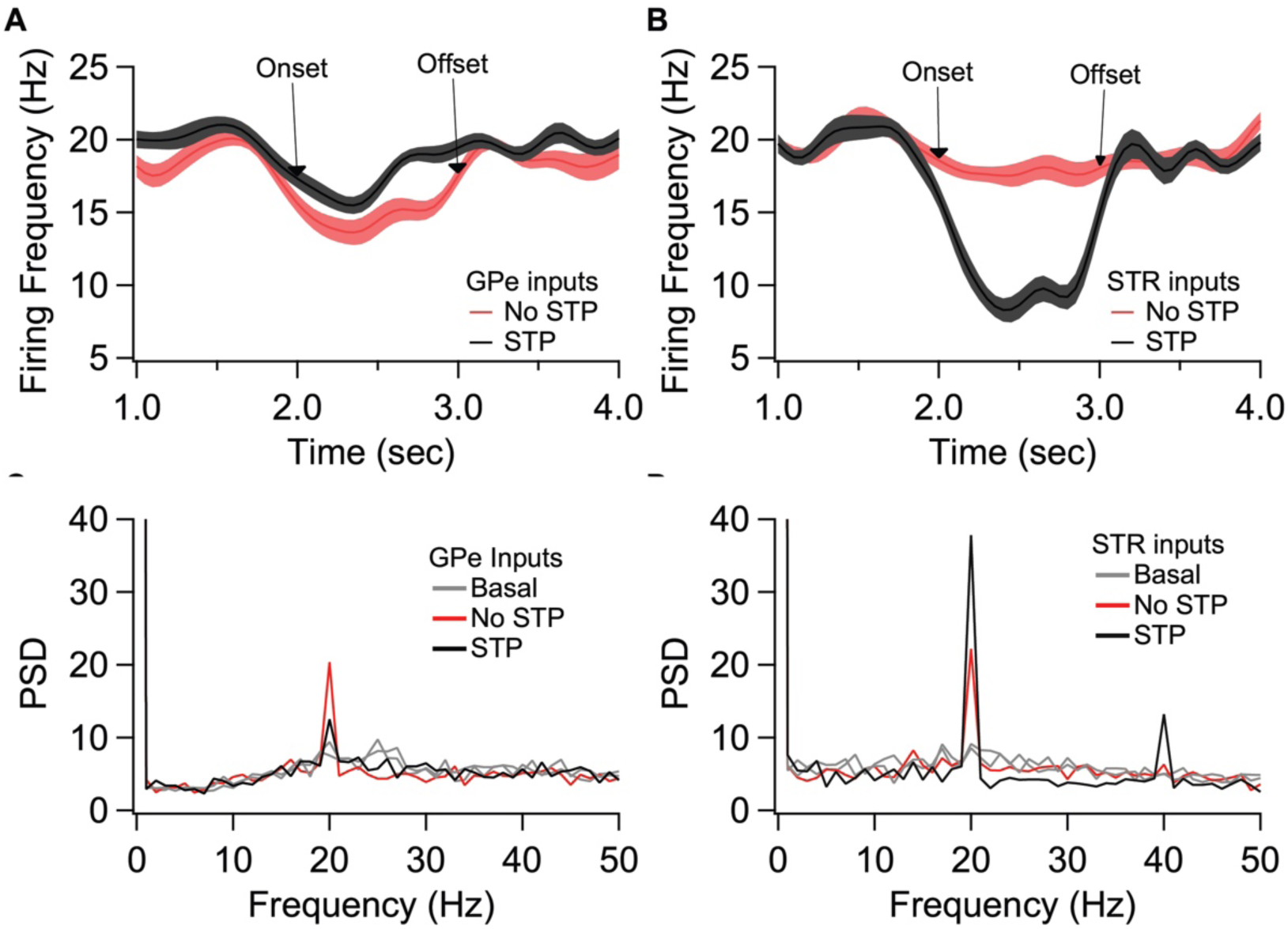
Effect of short term plasticity (STP) on EP firing frequency and phase locking. EP neurons were simulated in response to log normally distributed GPe (18 Hz), SPN (4 Hz) and STN (18 Hz) synaptic input, with an additional 20 Hz train from either GPe (A1, B1) or STR (A2,B2) delivered between 2 and 3 sec (marked by arrows labeled onset and offset). (A) Effect of STP on EP firing frequency. STP reduced the effect of 20 Hz GPe inputs on EP firing and enhanced the effect of 20 Hz STR inputs on EP firing. Shading shows +/− 1 SEM. (B) Amplitude of the PSD peak at 20 Hz (PSD calculated from the data segment between 2 and 3 sec) shows that STP reduced phase locking to GPe inputs but increased phase locking to STR inputs. Basal shows the PSD measured from the same simulations using data segments from 1-2 and 3-4 sec.

We then evaluated the effect of long-term plasticity of STR and GPe inputs. We simulated the response to STR, GPe and STN synaptic inputs under three different conditions: Control, synaptic strength resulting from the post-HFS protocol, and synaptic strength resulting from the post-HFS protocol with dopamine blocked. Mean firing frequency of these synaptic inputs was similar to that measured experimentally (Kita and Kita, 2011; Wilson and Bevan, 2011), thus STR inputs fired at 4 Hz, GPe inputs at 29 Hz, and STN inputs at 18 Hz. To demonstrate how synaptic plasticity changes information transmission, we modulated the mean firing frequency (using an inhomogeneous Poisson process), with a different oscillation frequency for STN, GPe and STR (Fig 9C). The information transmitted by the EP is represented as the amplitude of the PSD at the oscillation frequency for each structure. Fig 9A shows that synaptic plasticity (Post-HFS) increases the STR (direct pathway) information while reducing the GPe and STN information, but this enhancement is eliminated with dopamine blocked. We also performed simulations using log-normally distributed synaptic inputs, which captures the long tailed inter-spike-interval distributions observed in vivo (Kita and Kita, 2011). Fig 9B shows that the post-HFS protocol decreases energy at 20 Hz (β frequency), whereas post-HFS with dopamine blocked has enhanced energy at 20 Hz.

**Figure 9:**
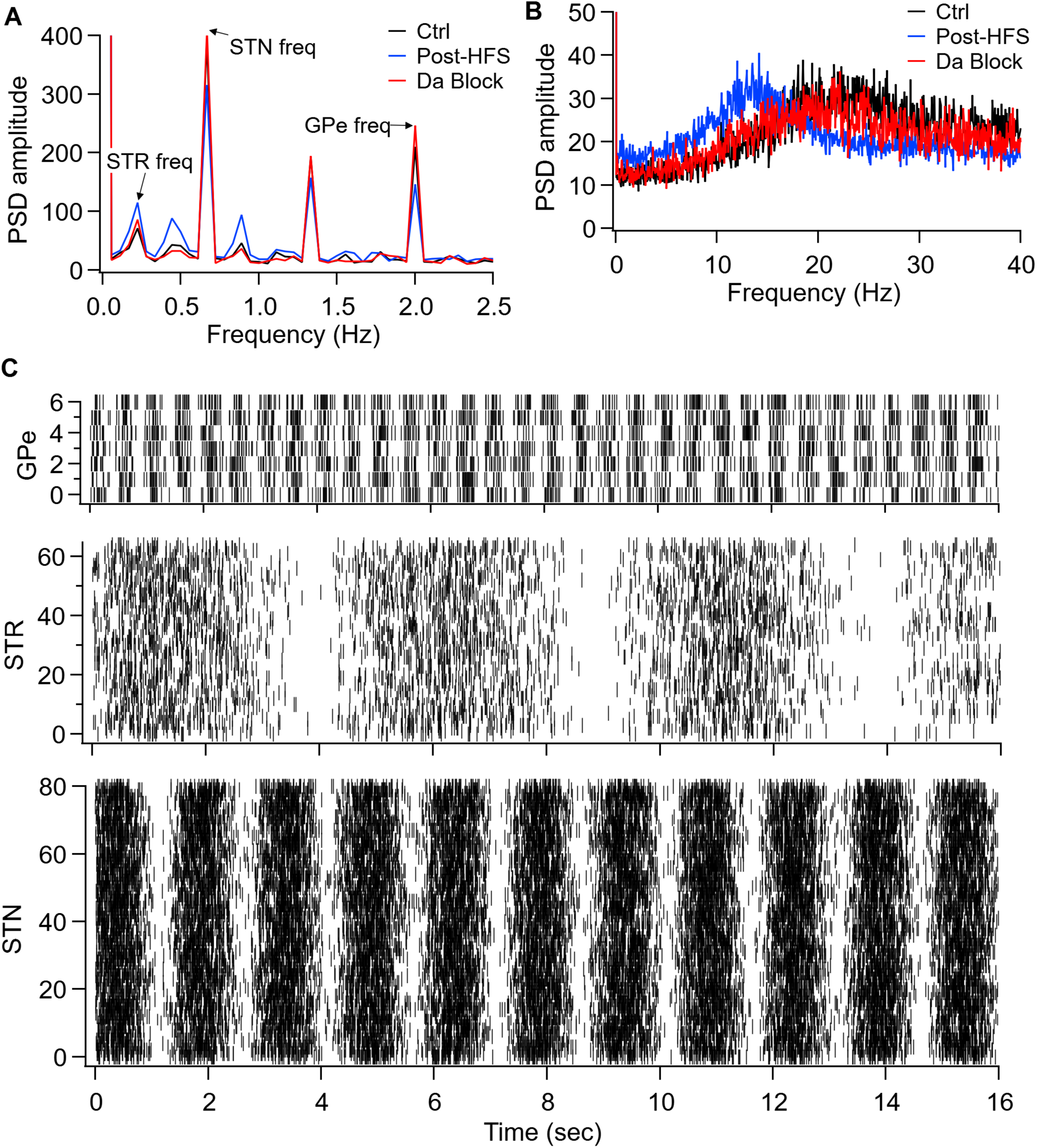
Model simulations demonstrate the eCB dependent synaptic plasticity enhances information transmission from the striatum and reduces energy in the β band. (A) Simulations in response to oscillatory synaptic inputs reveals that Post-HFS enhances energy at the STR frequency while reducing energy at GPe and STN frequencies. N=15 trials. (B) Simulations in response to log normally distributed synaptic inputs demonstrates that Post-HFS reduces β-band energy, whereas blocking dopamine increases energy in the β-band. N=15 trials. (C) Raster plots of synaptic inputs for 1 trial of simulations in (A). Oscillation frequency of 0.2 Hz (STR), 0.6 Hz (STN) and 2 Hz (GPe) modulates the mean firing frequency of synaptic inputs. Note that mean firing frequency of synaptic inputs was 4 Hz (STR), 18 Hz (STN) and 29.3 Hz (GPe) for both oscillatory (A) and log normally distributed (B) inputs.

## Discussion

Here we present results supporting a new integration role for dopamine and eCBs in modulating BG output. Using electrophysiology, optogenetics, and numerical modeling, we presented evidence suggesting interdependence between the direct, indirect, and hyperdirect afferents to the EP. We demonstrated that eCB modulates the synaptic strength of the indirect pathway (Fig. 1), the direct pathway (Fig, 2), and glutamatergic (Fig. 3) inputs to the EP, and that dopamine modulates eCB induced plasticity of all inputs to the EP (Fig. 5). We also showed that eCBs diffuse, modulating synaptic strength of neighboring EP neurons (Fig. 4). Thus, despite the lack of axon collaterals, information is transferred between neurons in the EP via endocannabinoid diffusion. Our numerical simulations demonstrate that the combination of STP and eCB mediated long-term plasticity enhances the transmission of direct pathway information while reducing indirect and hyper-direct pathway information. Thus, our results propose that the passage of BG information through the bottleneck of the EP allows dramatic modulation of the information using small neuronal populations. Our data suggest that this cellular mechanism controls the balance between direct and indirect pathway dominance of EP output.

eCBs are involved in neural functions ranging from homeostasis to cognition (Katona and Freund, 2012). Behavioral adaptations rely on changes in synaptic strength and the prevalence of eCB-mediated long-term depression (eCB-LTD) at synapses (Castillo et al., 2012). eCBs have been identified as triggers for short and long-term plasticity at synapses throughout the brain. There is a notable expression of CB1Rs in the basal ganglia, mainly in the striatum, GP, EP, and SNr (Herkenham et al., 1990). Activation of the CB1R modulates LTD and also short-term plasticity (Kreitzer and Regehr, 2001; Llano et al., 1991; Morishita and Alger, 2000; Ohno-Shosaku et al., 2001; Pitler and Alger, 1992; Wilson and Nicoll, 2001). This modulation of short-term plasticity appears in several basal ganglia regions including the striatum (Centonze et al., 2007; Freiman et al., 2006; Kreitzer and Malenka, 2005; Maccarrone et al., 2008; Narushima et al., 2007, 2006a, 2006b; Uchigashima et al., 2007; Yin and Lovinger, 2006), GP (Engler, 2005), SNr (Engler, 2005; Szabo et al., 2006; Yanovsky et al., 2003), SNc (Szabo et al., 2006) and NAc (Rancz and Ha, 2006). Furthermore, eCB induced LTD has been reported in several brain regions such as the dorsal striatum (Kreitzer and Malenka, 2007, 2005; Ronesi and Lovinger, 2005; Wang et al., 2006), NAc (Mato et al., 2008; Robbe et al., 2002), the cortex (Lafourcade et al., 2007; Nevian and Sakmann, 2006; Panikashvili et al., 2005), the cerebellum (Tzounopoulos et al., 2007), amygdala (Azad et al., 2004; Marsicano et al., 2002) and the hippocampus (Chevaleyre et al., 2007; Chevaleyre and Castillo, 2004, 2003). Here we demonstrated that glutamatergic input to the EP exhibit eCB-LTD that is induced by postsynaptic firing. HFS of corticostriatal glutamatergic inputs to SPNs is known to induce LTD that requires postsynaptic Ca^2+^ elevation and leads to a decrease in the probability of glutamate release (Calabresi et al., 1994, 1992; Choi and Lovinger, 1997; Kreitzer and Malenka, 2005). Striatal LTD is dependent on group I mGluRs and L-type Ca^2+^ channels. Ca^2+^ elevation by an LTD induction protocol induces eCB release. The eCB induced LTD reported here shares similarities with striatal LTD. The depression level of the STN-EP synapses depends on the frequency of the APs of the EP neurons. This matches the AP frequency dependence of Ca^2+^ concentration in the soma and the dendrite of the EP neurons that we demonstrated previously (Gorodetski et al., 2018), suggesting that the observed eCB-LTD is indeed calcium-dependent. Postsynaptic release of eCBs is sufficient to induce CB1R-mediated depression at GABAergic synapses (Adermark et al., 2009). Similarly, the synaptic depression reported here is blocked by AM-251, implicating CB1R in this LTD. On the other hand, in the direct pathway, STR-EP synapses exhibit eCB-LTP that also depends on the frequency of the APs and is blocked by CB1Rs antagonist. An eCB-LTP also has been observed in the dorsal striatum (Cui et al., 2015).

After identifying the eCBs effect on the EP pathways, we demonstrated that dopamine modulated the eCB induced plasticity (Fig. 5). Striatal evoked IPSCs display short-term facilitation modulated by dopamine via D1LR. Furthermore, GP evoked IPSCs that show short-term depression, are modulated by dopamine via D2LR (Lavian et al., 2017; Lavian and Korngreen, 2016). Moreover, D1R is co-localized with striatal axon terminals, and D2R and D3R are co-localized with GP axon terminals (Lavian et al., 2018). In the current study, D2 and D1 antagonists blocked the eCB effects in the STN-EP and Str-EP synapses, respectively. These results indicate that DA modulates the generation of eCB-LTD by synaptic activity and comply with previous work implicating dopamine modulation of state-dependent eCB release in the striatum (Kreitzer and Malenka, 2007, 2005). However, in the GP-EP synapses, the block of D2R resulted in LTP instead of LTD. These results indicate that the LTD or LTP depends on the activation of D2Rs and CB1Rs, which differs from corticostriatal synapses (Xu et al., 2018), where eCB-LTP requires D2Rs. Using optogenetic stimulation, we showed that dopamine is being released by electrical stimulation, as dopamine agonists had to be included to produce LTP in the absence of electrical stimulation. Finally, we demonstrate that eCB diffuses and modulate synaptic plasticity in neighboring neurons (Fig. 4). The diffusion of eCB implies that all three types of synapses likely undergo plasticity at the same time. Moreover, nearby neurons that do not fire at high frequency may experience modulation of synaptic inputs. These results suggest that the EP is not only a feedforward nucleus (Parent et al., 2000), although it contains no axon collaterals.

We created a data-driven model of EP neurons to evaluate the functional effect of the simultaneous plasticity of all three synapses. We measured the response to in vivo like inputs to STR, STN and GPe synapses simultaneously. Our modeling results show that eCB mediated long term plasticity enhances information transmission from the direct pathway, compared to indirect and hyper-direct pathways, and the enhanced direct pathway information is eliminated when dopamine is blocked, showing the role of dopamine in controlling basal ganglia output. Many models of the basal ganglia have evaluated the role of dopamine in action selection. In most of these models (Humphries et al., 2006; Leblois et al., 2006; Lindahl and Kotaleski, 2016), dopamine influences direct vs indirect pathway information by its effect on striatal activity – either excitability or corticostriatal synaptic plasticity. One model implemented short term depression of GPe and STN inputs to SNr (another BG output region), together with dopamine modulation of STR to GPe and STR to SNr synapses (Lindahl and Kotaleski, 2016). However, the contribution SNr input modulation to action selection was not evaluated. Several network models that included GPi neurons (primate analogue of EP) evaluated the effect of dopamine on GPi oscillatory firing, to investigate mechanisms underlying Parkinson’s beta oscillations and normalization by DBS (Hahn and McIntyre, 2010; Humphries and Gurney, 2012; Kumaravelu et al., 2016). In these models, dopamine modulates striatal or GPe neurons or synapses, and beta oscillations in the GPi are a readout of basal ganglia network state. In contrast, our model revealed a direct effect of dopamine on beta oscillations. Specifically, the model exhibits a small increase in power at beta frequency, produced solely by dopamine mediated changes in synaptic plasticity of EP neurons. Thus, this suggests that lack of dopamine in the GPi can contribute to the production of beta oscillations in Parkinson’s.

We can formulate a conceptual model for synaptic integration in EP neurons based on the anatomical and functional polarity of EP neurons. In this model, GABAergic inputs from the GP act, due to their short-term depression kinetics and high baseline firing rates, as somatic shunting inhibition delivering an almost constant inhibition to the soma (Bugaysen et al., 2013; Lavian and Korngreen, 2019). The spontaneous firing rate of striatal projection neurons (SPNs) is low; thus, synaptic input to EP dendrites will not be amplified by short-term facilitation. However, striatal activation by cortical and thalamic inputs, leading to a burst of SPN activity, will lead to amplification of dendritic inputs in the EP, phase locking of EP firing to that of the STR (Lavian et al., 2017; Lavian and Korngreen, 2016). Thus, short-term facilitation amplifies trains of GABAergic inputs from STR to dendrites of EP neurons, overcomes the somatic shunt, and allows short bursts of STR firing to affect the firing of neurons in the EP. It is possible to visualize GABAergic neurons in the EP as having a somatic obstacle surmounted by a dendritic amplifier.

The significant finding we presented here is that eCBs modulate all synaptic inputs to the EP. Thus, two variables control the inputs to the EP, the levels of eCBs, and dopamine. Since the level of eCBs is a function of the post-synaptic firing rate (Figs 1-3), it is possible to state that two state variables, dopamine release and post-synaptic firing rate, bind all synaptic inputs to the EP. Thus, our results suggest crosstalk between direct, indirect, and hyperdirect inputs to the EP. These inputs are not independent; global eCB modulation binds them together. It is fascinating to interpret these results using the conceptual model we presented above. In the presence of dopamine, an increase in EP firing (probably due to glutamatergic input from the STN) lowers the somatic barrier (eCB-LTD in GP-EP synapse) while boosting the dendritic amplifier (eCB-LTP in STR-EP synapse); increasing the weight of input from the direct pathway while lowering that from the indirect pathway (Fig. 10).

**Fig. 10.**
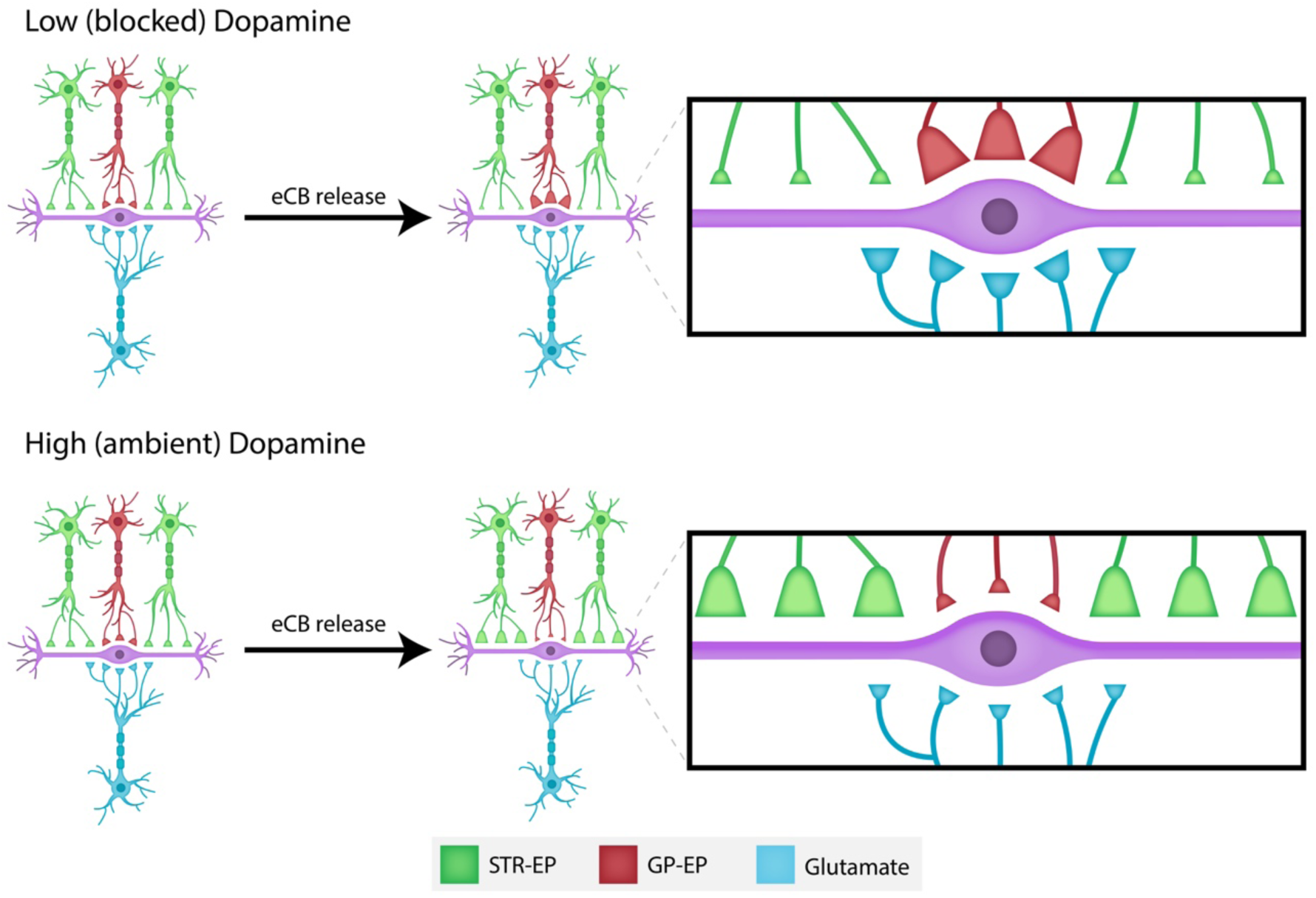
Schematic drawing of the dopamine-eCB induced synaptic dynamics in the EP.

Moreover, increased EP firing lowers the weight of glutamatergic input to the EP, further increasing the potentiated input from the striatum. In this conjecture, it is essential to note that neurons in the EP fire spontaneously. Therefore, we may expect a constitutive level of eCB release leading to maintained modulation of all inputs to the EP. When dopamine levels in the EP are low, the opposite happens. The somatic barrier is potentiated, and the dendritic amplifier muted (Fig. 10). An increase in EP firing releases eCBs that may, via diffusion, modulate neighboring synapses on the same neuron. This spatial effect may lower the somatic barrier even more while boosting the dendritic amplifier. We can perhaps visualize somatic inhibition as a dopamine-eCB modulated sluice gate, dendritic inhibition as a dopamine-eCB tunable amplifier, and glutamatergic excitation as being retrogradely dampened by eCBs. Furthermore, it is easy to envision the contribution of this cellular mechanism for *selecting* which information from the direct or indirect pathway, will dominate EP output. These results transform the prevailing view of the entopeduncular nucleus as a feedforward “relay” nucleus to an intricate control unit, which may play a vital role in the process of action selection.

## Acknowledgments

This work was supported by a grant from the Israel Science Foundation (#168/16).

## Author contribution

The authors designed the study, performed the experiments, analyzed data drafted and revised the manuscript. The authors read and approved the final version of the manuscript for publication.

## Data accessibility

Data will be available upon a request to corresponding author.

## Abbreviations

STN: subthalamic nucleus
EP: entopeduncular nucleus
SNr: substantia nigra reticulate
LTD: long-term depression
CB1: endocannabinoid 1
GP: globus pallidus
ACSF: artificial cerebrospinal fluid
DMSO: Dimethyl sulfoxide
ROI: regions of interest
F: Fluorescence
LTP: long term potentiation
AP: Action potential
SPNs: spiny projection neurons

